# DNA cytosine methylation modulates UV resistance and nucleotide excision repair gene expression in *Escherichia coli*

**DOI:** 10.64898/2026.06.22.733644

**Authors:** Shunsuke Ichikawa, Mika Okazaki

## Abstract

Bacterial survival after ultraviolet (UV) exposure is shaped not only by the extent of DNA damage but also by the physiological state-dependent capacity for DNA repair. Here, we examined the mechanisms underlying growth phase-dependent UV resistance in *Escherichia coli* K-12 exposed to 262 nm UV irradiation. Stationary-phase cells required higher UV fluence for log inactivation than exponential-phase cells, whereas the levels of UV-induced DNA damage, assessed by cyclobutane pyrimidine dimer staining and real-time PCR, did not differ markedly between the two growth phases. Deletion of nucleotide excision repair (NER) genes, including *uvrA*, *uvrB*, *uvrC*, and *uvrD*, markedly reduced survival after UV irradiation, indicating that NER is essential for the high UV resistance of stationary-phase cells. Quantitative real-time reverse transcription PCR showed stronger UV-induced expression of several DNA repair and UV resistance genes, including *uvrA*, *uvrB*, *cho*, *umuC*, and *umuD*, in stationary-phase cells than in exponential-phase cells.

Furthermore, deletion of the DNA cytosine methyltransferase gene *dcm* increased UV resistance and enhanced the expression of *uvrB*, *cho*, *umuC*, *umuD*, and *sulA* in stationary-phase cells. These findings suggest that DNA cytosine methylation modulates UV resistance in *E. coli*, at least in part by influencing NER- and SOS-associated gene expression.

## 1. Introduction

Continuous contamination of aquatic environments with pathogenic microorganisms exacerbates the incidence of serious human illnesses and requires the development of efficient disinfection methods [1, 2]. Ultraviolet (UV) radiation disinfection is a straightforward and effective method against a range of pathogens, and does not produce any byproducts. Mercury lamps with wavelengths of 253.7 nm are commonly used as UV light sources in water purification plants [3, 4]. Ultraviolet light-emitting diodes (UV-LEDs) represent an alternative light source that possess several benefits, including wavelength diversity, rapid start-up times, long lifetimes, comparatively low energy consumption, compactness, resilience, environmental friendliness (as they do not use mercury), and high-frequency on/off capabilities. Therefore, UV-LEDs have the potential to replace mercury UV lamps [5].

UV fluences for log inactivation by UV-LED have often been determined, and these conditions are required to design an appropriate UV-LED apparatus for water disinfection. However, the determined UV fluences for the log inactivation of a model bacterium *Escherichia coli* by UV-LED are not identical among previous studies that reported that 6-10.8 mJ/cm^2^ by 265 nm UV-LED and 9-13.8 mJ/cm^2^ by 280 nm UV-LED were required for 4 log inactivation [6, 7]. We recently reported that the inactivation of *E. coli* stationary phase cells requires a higher UV fluence compared to that for exponential phase cells [8]. Similar phenomena have been reported for other bacteria, including *Cronobacter sakazakii*, *Staphylococcus aureus*, *Salmonella typhimurium* [9–11]. From these reports, it is expected that bacterial UV resistance fluctuates depending on the growth conditions.

UV-C radiation induces DNA lesions such as cyclobutane pyrimidine dimers (CPD) and (6-4) photoproducts in bacterial cells [12]. These DNA lesions inhibit the replication and transcription of DNA, ultimately resulting in cell death. The bactericidal impact is contingent not only on the UV-C dosage but also on the bacterial capacity to repair DNA damage [13]. Nucleotide excision repair (NER) is a major DNA repair system that occurs in both prokaryotes and eukaryotes. DNA damage by UV irradiation activates RecA proteins that induce the SOS response and promote gene expression of DNA repair systems, including NER [14]. The molecular mechanisms underlying NER in *E. coli* are well understood. The UvrA-UvrB complex monitors DNA and the UvrA subunit recognizes distortions in the double-helix structure. UvrA is released when the UvrA-UvrB complex binds to DNA damage. UvrB dissociates base pairs between DNA strands near DNA damage sites. UvrB then recruits UvrC, which generates nicks on the damaged DNA strand. UvrD releases damaged DNA strands. Finally, the gap is repaired using DNA polymerase and ligase [14]. Other SOS response genes, *umuC* and *umuD*, specifically inhibit the transition from stationary phase to exponential growth, and this correlates with increased survival after UV irradiation [15]. Another SOS gene, *sulA*, is also a UV-inducible cell division inhibitor [16].

UV-induced DNA damage is also repaired by photolyase activity that is a product of the *phr* gene. Photolyase converts thymine dimers (a major type of CPD) into two normal thymines with blue light energy [17]. The induction of *phr* gene expression was independent of the SOS response.

In this study, we demonstrated the fluctuation of the required UV fluence for *E. coli* inactivation using a dose-response curve. Subsequently, we demonstrated that DNA repair systems, including NER, were highly induced by UV irradiation in UV-resistant *E. coli* stationary phase cells. Furthermore, we revealed that DNA cytosine methylation affects NER induction in *E. coli*.

## 2. Materials and methods

### 2.1 Inactivation of *E. coli* cells using UV-LED

A single-gene knockout Keio mutant collection of *E. coli* K-12 was obtained from the National BioResource Project-*E. coli* of the National Institute of Genetics in Japan. The *E. coli* strains were cultured aerobically in Luria-Bertani medium at 37 °C. When an OD_600_ of 0.3 was reached, the exponential phase cells were harvested by centrifugation at 5,000 × *g* for 3 min. Alternatively, *E. coli* was cultured for 16 h, and harvested as the stationary phase cells. The cells were then washed and resuspended in phosphate-buffered saline (PBS). An *E. coli* cell suspension (10^6^ or 10^8^ CFU/mL) was transferred to a 5.5 cm-diameter glass Petri dish to provide a sample depth of 8.0 mm. A 262 nm UV-LED (Nikkiso Giken Co. Ltd., Ishikawa, Japan) was positioned 32 mm above the surface of the *E. coli* suspension. During the UV irradiation experiment, the *E. coli* suspension was stirred at 700 rpm. The average irradiance at the surface of the *E. coli* suspension was 0.242 mW/cm^2^.

The UV-irradiated *E. coli* cells were cultivated in a Petrifilm aerobic count plate (3M, MN, USA) in the dark at 35 °C for 48 h after being exposed to 262 nm UV for specific exposure periods of 0-50 s. Colony-forming units (CFUs) were used to determine the number of viable cells in 1 mL of *E. coli* suspension. The viable *E. coli* cells (CFU/mL) in the UV-treated samples (N) were standardized to those in the untreated samples (N_0_). The log-linear component of the fluence-response curve was used to calculate the inactivation rate constant for *E. coli* inactivation using a 262 nm UV-LED [3, 18].

### 2.2 Evaluation of DNA damage by UV irradiation

CPD in UV-treated *E. coli* cells was detected using the OxiSelect™ Cellular UV-Induced DNA Damage Staining Kit (CPD) (Cell Biolabs, CA, USA) according to the manufacturer’s instructions. Briefly, UV-treated *E. coli* cells (10^8^ CFU) were fixed in 75% methanol/25% acetic acid and 70% ethanol. After blocking, CPDs were detected using an anti-CPD antibody and FITC-labeled secondary antibody. Bacstain 4′,6-diamidino-2-phenylindole (DAPI) solution (Dojindo, Kumamoto, Japan) was used to stain total DNA in the bacterial cells. The stained *E. coli* cells were observed under fluorescent microscopes, BZ-X800 (Keyence, Osaka, Japan) and BX-51 (Olympus, Tokyo, Japan).

DNA damage caused by UV irradiation has also been evaluated using real-time PCR [19]. Genomic DNA was extracted from 6 mJ/cm^2^ UV-irradiated *E. coli* cells using a Wizard^®^ Genomic DNA Purification Kit (Promega, WI, USA). The primers were designed using Primer3 software (Table S1). Each 20 μL reaction mixture contained a DNA template, 0.5 µM of each primer, and PowerUp SYBR Green Master Mix (Thermo Fisher Scientific, MA, USA). A StepOnePlus Real-Time PCR System (Thermo Fisher Scientific) was used to conduct real-time PCR with triplicate measurements. Analyses of the melt curves confirmed the primer annealing specificity and ensured the absence of a secondary structure.

### 2.3 Gene expression analysis by quantitative real-time RT-PCR

*E. coli* cells were inactivated by 6 mJ/cm^2^ 262 nm UV irradiation. After 30 min, the cells were lysed, and total RNA was extracted using MN Bead Tubes Type B Plus (Macherey-Nagel, Düren, Germany) and NucleoSpin^®^ RNA Plus (Macherey-Nagel, Düren, Germany) according to the manufacturer’s instructions. Reverse transcription of RNA was conducted according to the manufacturer’s instructions using a ReverTra Ace qPCR RT Master Mix with gDNA Remover kit (TOYOBO, Osaka, Japan). The real-time RT-PCR was conducted as described in Section 2.2. The data were standardized to the level of *gapA* expression [20].

## 3. Results

### 3.1 Inactivation of *E. coli* exponential and stationary phase cells by 262 nm UV irradiation

*E. coli* cells in the exponential and stationary phases were inactivated by 262 nm UV-LED irradiation (Fig. 1). The 1 log (90%) inactivation of stationary phase cells required 2.5-fold higher 262 nm UV fluence compared to that of exponential phase cells (Table 1). The required fluences for 3 log and 4 log inactivation of the exponential and stationary phase cells were 6.8 and 8.7 mJ/cm^2^ and 8.4 and 9.8 mJ/cm^2^, respectively.

**Fig. 1.**
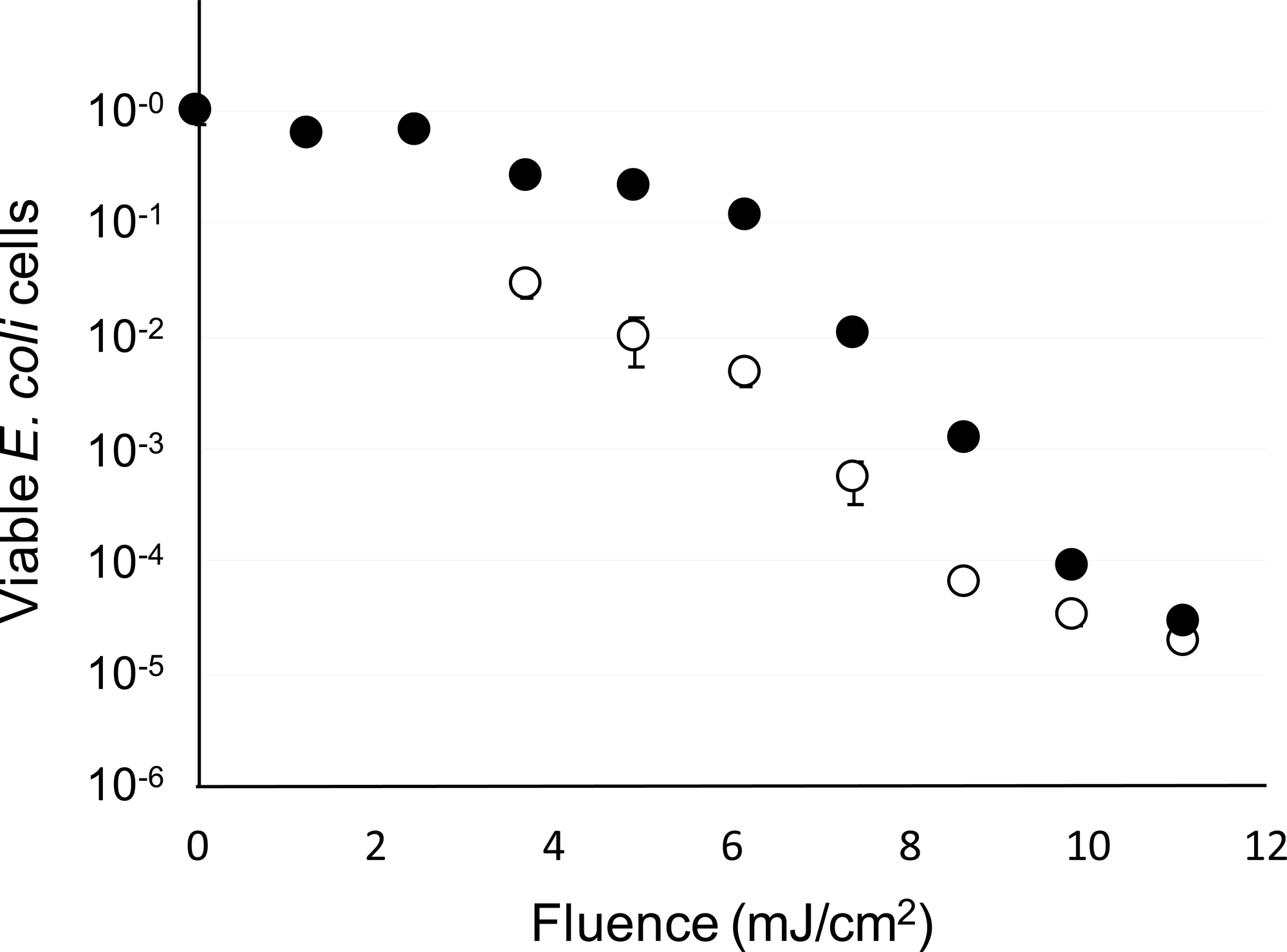
UV-LED inactivation of *E. coli* exponential and stationary phase cells by 262 nm irradiation. *E. coli* K-12 (10^6^ CFU/mL) cells from the exponential phase (white dots) and stationary phase (black dots) were inactivated by 262 nm UV-LED. Error bars indicate standard error.

### 3.2 DNA damage in *E. coli* exponential and stationary phase cells after the UV irradiation

CPD generated by UV irradiation was successfully detected in UV-treated *E. coli* cells but not in untreated cells. No significant difference in FITC signal intensity between the exponential and stationary phase cells was observed (Fig. 2a). DNA damage caused by UV irradiation was also evaluated using real-time PCR [19]. The quantity of detected DNA was decreased in response to UV treatment; however, there were no significant differences between exponential and stationary phase cells (Fig. 2b). These results suggest that the level of DNA damage induced by UV irradiation was not significantly different between exponential and stationary phase cells.

**Fig. 2.**
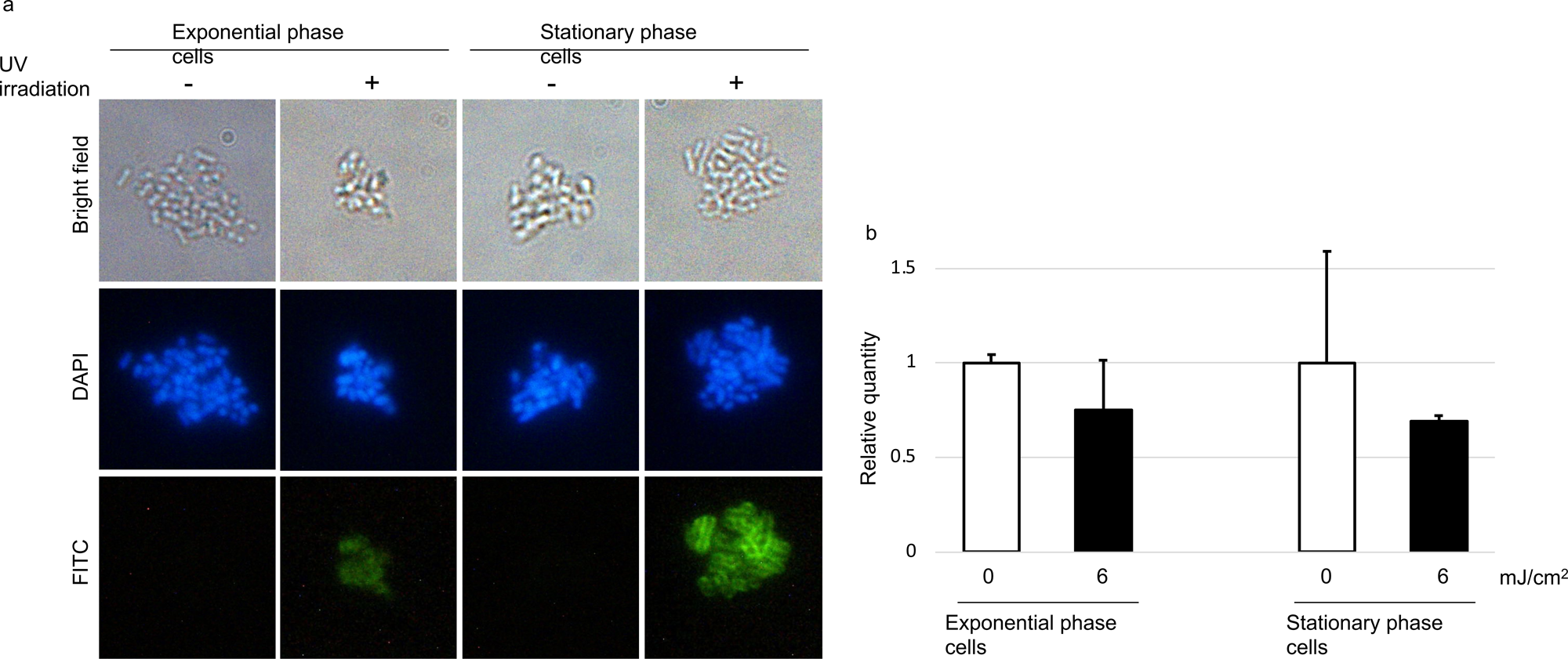
DNA damage induced by 262 nm UV inactivation in *E. coli*. (a) CPDs in UV-treated *E. coli* cells were detected with anti-CPD antibody and FITC-labeled secondary antibody that is presented in green. DNA was stained with DAPI as indicated by blue staining. (b) DNA damage in response to UV irradiation was evaluated by real-time PCR. Measurements were performed in triplicate. Error bars indicate standard error. The levels of DNA damage are equal between the exponential and stationary phase cells.

### 3.3 Inactivation of *E. coli* nucleotide excision repair gene mutants by 262 nm UV irradiation

*E. coli* stationary phase cells (10^8^ CFU/mL) were inactivated to a level of 1.3 x 10^-1^ by 6 mJ/cm^2^ 262 nm UV irradiation. In contrast, the *uvrA* deletion strain drastically reduced the viable cell number to 5.9 x 10^-7^ in response to UV irradiation (Fig. 3). Similarly, single deletion of the *uvrB*, *uvrC*, and *uvrD* genes significantly reduced the survival fractions after UV irradiation to 1.3 x 10^-5^, 8.7 x 10^-5^, and 1.2 x 10^-3^, respectively. The deletion mutant strain for *recA*, which is a master regulator gene for the SOS response, also exhibited reduced resistance to UV irradiation. These results suggest that the NER system regulated by the SOS response is essential for high UV resistance in *E. coli* stationary phase cells. The SOS response in *E. coli* cells is induced by UV irradiation. We observed that the expression level of *uvrA* was significantly increased by 8.8-fold in the stationary phase after UV irradiation, while the expression level was only increased by 1.6-fold in the exponential phase cells (Fig. 4a). Similarly, the expression level of *uvrB* was significantly increased by 5.1-fold in the stationary phase after UV irradiation, while the expression level was only increased by 1.7-fold in the exponential phase cells. The expression of the *uvrC* homolog *cho* was drastically increased by 36.6-fold in stationary phase cells after UV irradiation but was only increased to 2.4-fold in exponential phase cells. Elevated expression levels of *umuC* and *umuD* in stationary phase cells were also observed after UV irradiation. These results suggest that genes related to UV resistance were strongly induced in stationary-phase cells after UV irradiation (Fig. 4a). The expression of *phr*, a key gene for photoreactivation after UV irradiation, is not regulated by the SOS response [21]. The expression level of *phr* was also higher in stationary phase cells than it was in exponential phase cells; however, this increase did not depend upon UV irradiation.

**Fig. 3.**
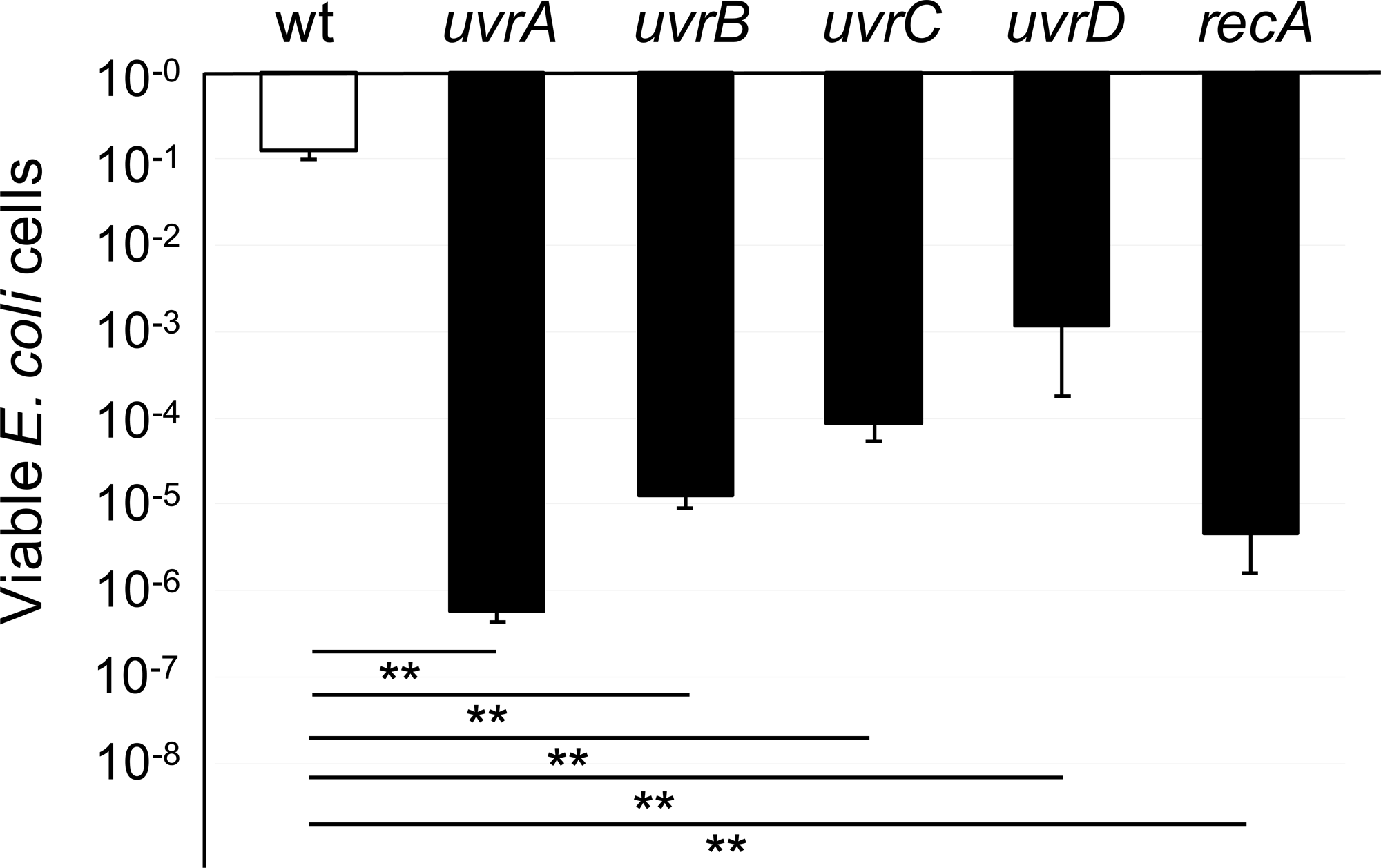
The loss of UV resistance in *E. coli* mutant strains. Stationary phase cells (10^8^ CFU/mL) of the *E. coli* mutant strains *uvrA*, *uvrB*, *uvrC*, *uvrD*, and *recA* were inactivated by 6 mJ/cm^2^ 262 nm UV irradiation. Error bars indicate standard error. Statistical significance was assessed using a two-tailed Student’s t-test. **p < 0.01. The experiments were performed in triplicate.

**Fig. 4.**
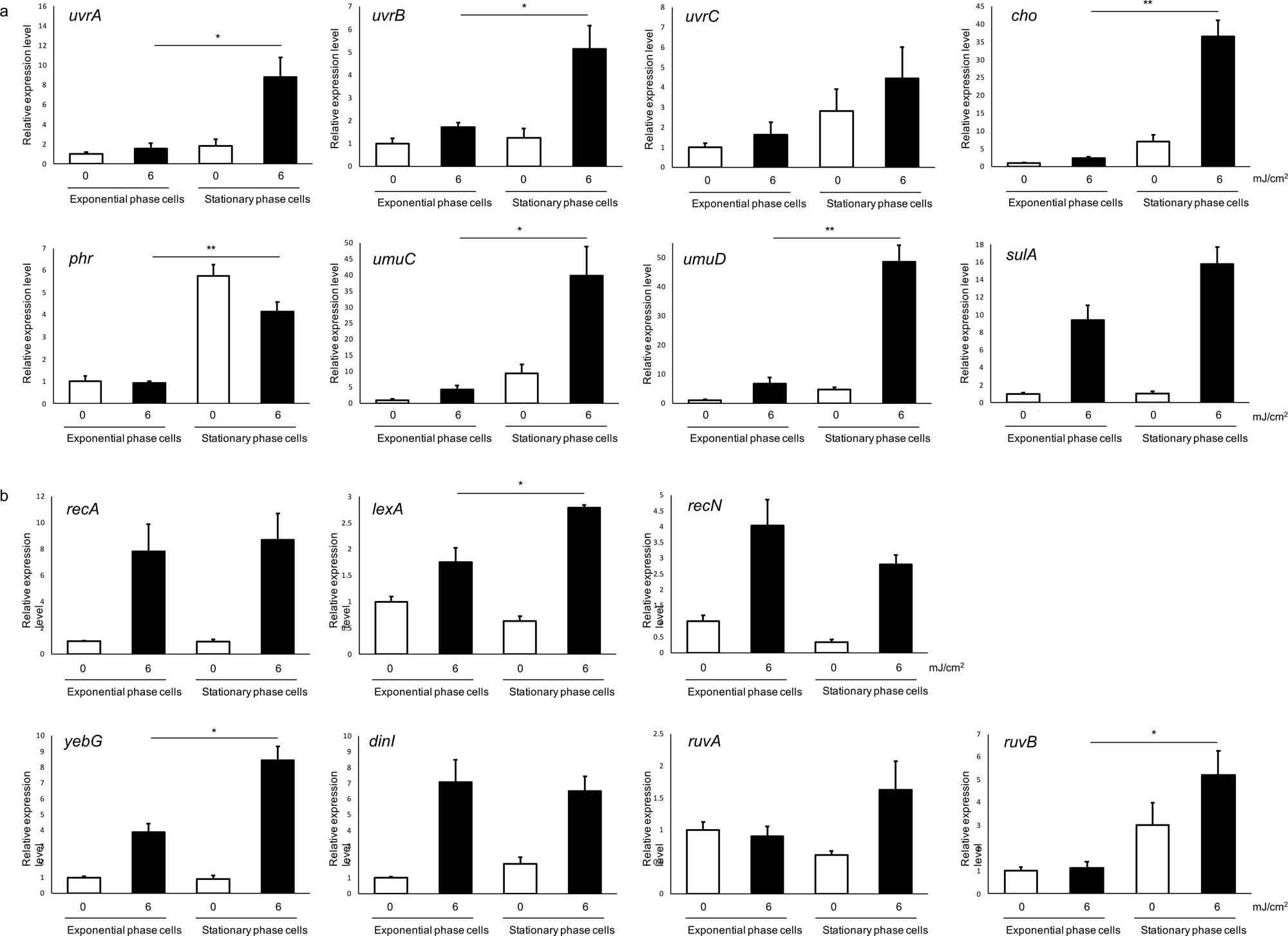
The gene expression levels in *E. coli* exponential and stationary phase cells after UV irradiation. The expression of (a) UV resistance genes and (b) other SOS genes in UV-treated *E. coli* cells was evaluated using real-time qPCR. Error bars indicate the standard error. Statistical significance was assessed using a two-tailed Student’s t-test. *p < 0.05, **p < 0.01. Measurements were performed in triplicate.

Subsequently, we evaluated the expression levels of other SOS response genes (Fig. 4b). The expression levels of *lexA*, *yebG*, and *ruvB* were also higher in stationary-phase cells after UV irradiation; however, the expression levels of *recA*, *recN*, *dinI*, and *ruvA* were not significantly different between exponential- and stationary-phase cells. These results suggest that the phase dependence of UV-induced SOS gene expression was gene-specific.

### 3.4 Inactivation of *E. coli* DNA methyltransferase gene mutants by 262 nm UV irradiation

The DNA adenine methyltransferase gene mutant *dam* and the DNA cytosine methyltransferase gene mutant *dcm* were inactivated by 6 mJ/cm^2^ 262 nm UV irradiation (Fig. 5). The viable *dam* and *dcm* exponential phase cells after UV irradiation were 1.3 x 10^-2^ and 1.4 x 10^-1^, respectively. The viable *dam* and *dcm* stationary phase cells after UV irradiation were 1.2 x 10^-1^ and 5.8 x 10^-1^, respectively. These results indicate that the *dcm* deletion mutant, in particular, exhibited higher UV resistance than the wild-type strain.

**Fig. 5.**
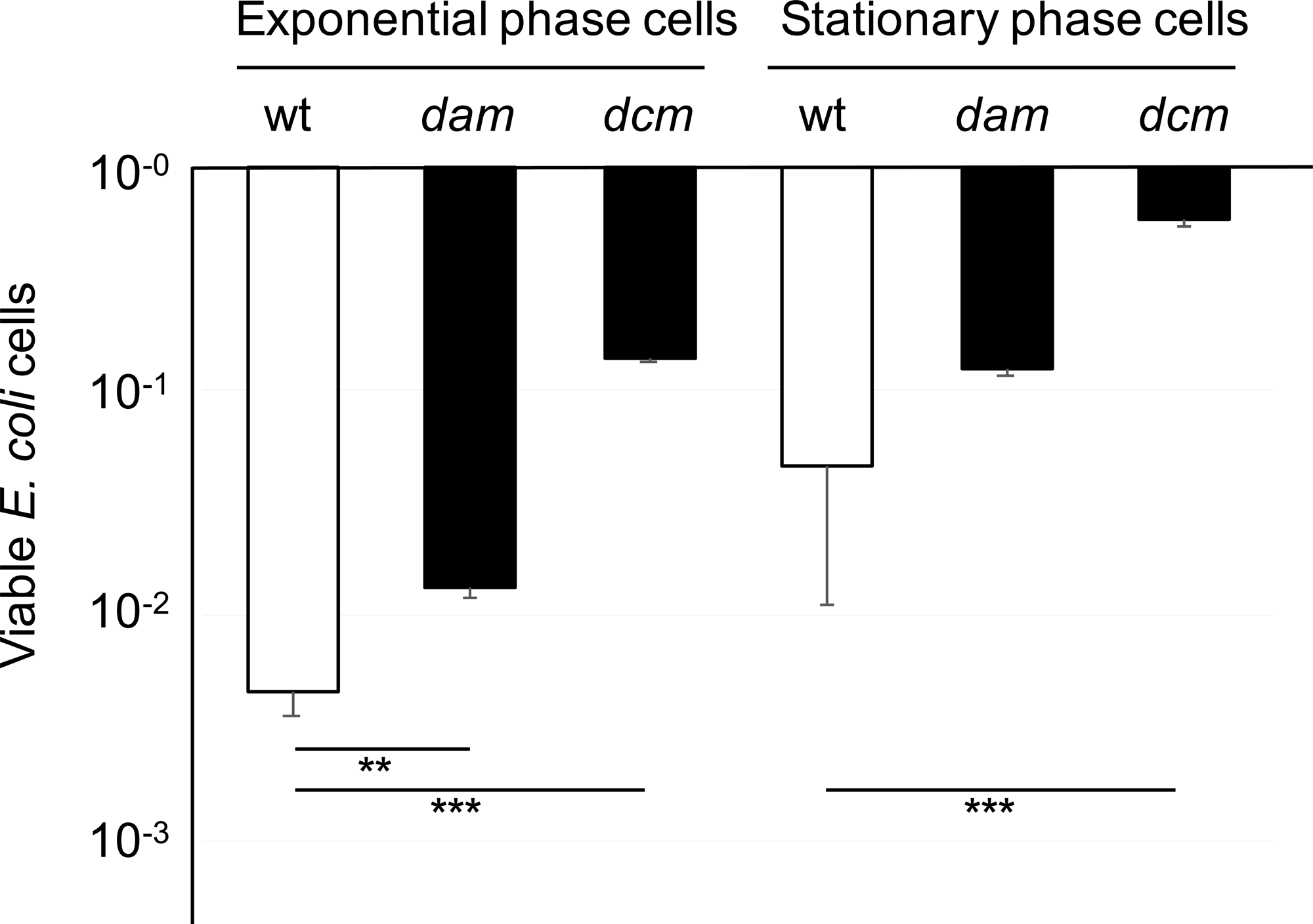
The UV resistance of *E. coli* DNA methyltransferase mutants. *E. coli dam* and *dcm* mutant strains were inactivated by 6 mJ/cm^2^ 262 nm UV irradiation. Error bars indicate standard error. Statistical significance was assessed using a two-tailed Student’s t-test. **p < 0.01, ***p < 0.001. The experiments were performed in triplicate.

The expression levels of UV resistance genes in wild-type and *dcm* stationary phase cells after UV irradiation were next evaluated (Fig. 6). The *uvrA* expression did not change between the wild-type and *dcm* strains, while *uvrB* and the *uvrC* homolog *cho* expression levels were increased in the *dcm* strain. The expression of other UV resistance genes (*umuC*, *umuD*, *and sulA*) was also enhanced in the *dcm* strain. The expression of other SOS response genes (*recA* and *lexA*) was also increased in *the dcm* strain. These results suggest that DNA cytosine methylation represses the induction of the SOS response, including the expression of the NER genes *uvrB* and *cho* and other UV resistance genes (*umuC*, *umuD*, *and sulA*).

**Fig. 6.**
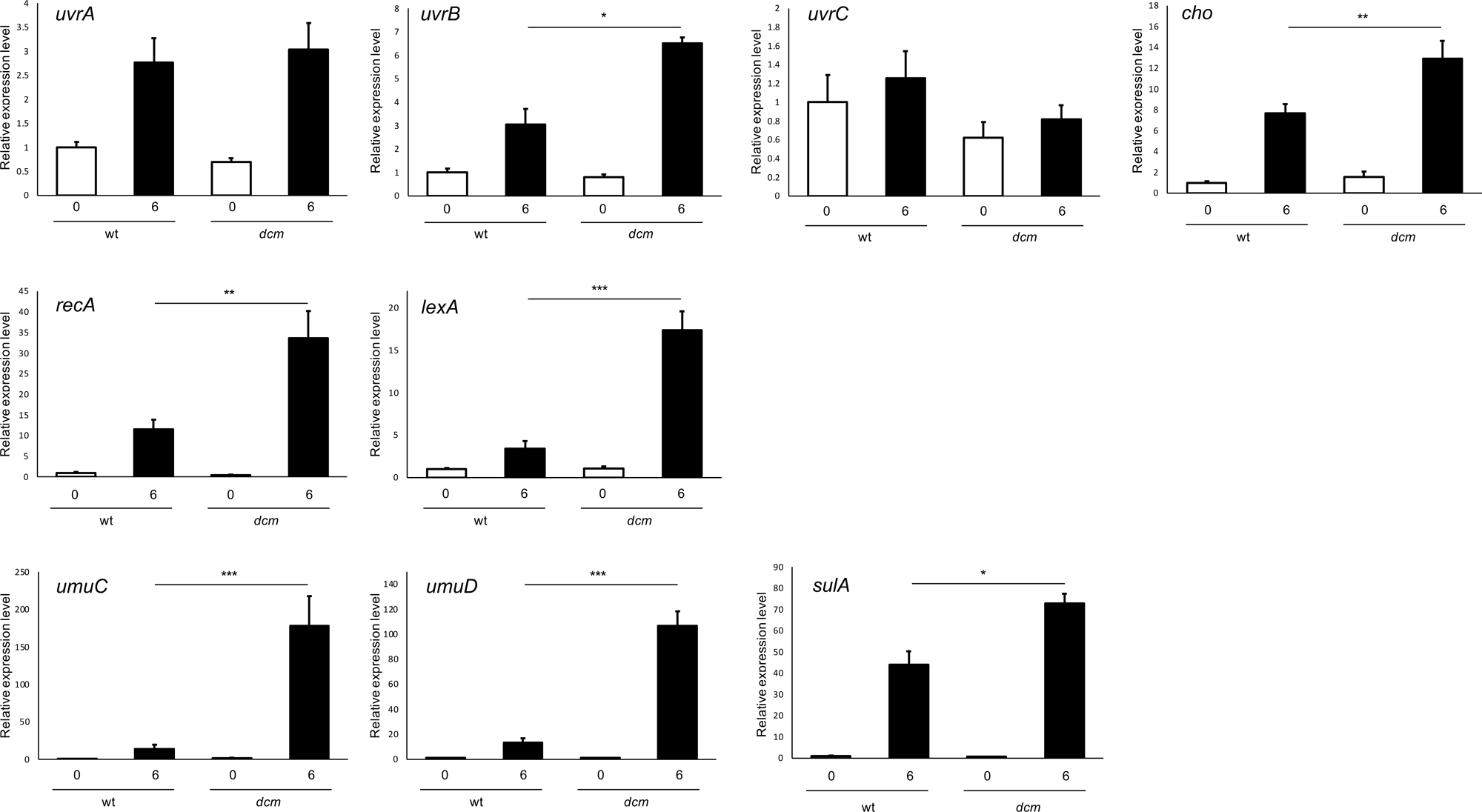
The gene expression levels in the *E. coli dcm* strain after UV irradiation. The expression of SOS genes in UV-treated *E. coli dcm* stationary phase cells was evaluated by real-time qPCR. Error bars indicate the standard error. Statistical significance was assessed using a two-tailed Student’s t-test. *p < 0.05, **p < 0.01, ***p < 0.001. Measurements were performed in triplicate.

## 4. Discussion

The required fluences were 8.4-9.8 mJ/cm^2^ for 4 log inactivation of *E. coli* cells by 262 nm UV irradiation, and these were consistent with values reported in previous studies [6, 7] (Fig. 1). We again demonstrated the high UV resistance of stationary-phase cells compared to that of exponential-phase cells as reported in other bacteria [8–11].

We next attempted to reveal the molecular mechanism underlying high UV resistance in *E. coli* stationary phase cells. No significant difference was observed in UV-induced DNA damage between the exponential and stationary phase cells (Fig. 2). The NER gene mutants *uvrA*, *uvrB*, *uvrC*, and *uvrD* lost their UV resistance (Fig. 3). The expression of DNA repair genes was induced in stationary-phase cells after UV irradiation (Fig. 4a). Thus, the DNA repair system strongly contributes to the UV resistance of *E. coli* stationary-phase cells.

The expression of the DNA repair genes *uvrA*, *uvrB*, and *cho* is regulated by the SOS response. As we mentioned previously, the expression of these genes was drastically induced in the stationary phase cells after UV irradiation; however, the expression of *recA*, a master regulator of the SOS response, was equally induced in both the exponential and stationary phase cells (Fig. 4b). Thus, it appears that other gene regulation systems also contribute to the expression of these UV resistance genes.

Recent studies have reported that DNA modifications play a role in gene expression regulation through a process known as epigenetic regulation not only in eukaryotes but also in prokaryotes. DNA adenine methylation has been identified as a gene expression regulator in bacteria that functions principally by altering the activities of transcription factors whose effects are associated with the methylation status of the target. It is reported that DNA adenine methylation is implicated in the pathogenicity of diverse bacteria such as *Salmonella enterica*, *Mycobacterium tuberculosis*, *Streptococcus mutans*, *Aggregatibacter actinomycetemcomitans*, *Vibrio cholerae*, *Aeromonas hydrophyla*, *Pasteurella multocida*, and enterohemorrhagic *E. coli* [22, 23]. As an example of bacterial epigenetic regulation, the expression of most adhesin genes is regulated by DNA adenine methylation patterns in *E. coli* [24]. However, only a few roles for cytosine methylation in the regulation of gene expression have been reported in bacteria. For example, Militello *et al*. reported that cytosine methylation regulates ribosomal gene expression during the stationary phase [25]. In bacterial genomic DNA, N^6^-methyl-adenine (6mA), N^4^-methyl-cytosine (4mC), and C^5^-methyl-cytosine (5mC) are formed at specific target sites [26, 27]. DNA adenine methyltransferase that binds to the GATC motif is the most thoroughly investigated single methyltransferase in *E. coli* [23]. DNA cytosine methyltransferase Dcm, the only cytosine methyltransferase in *E. coli*, methylates the internal cytosine of the CCWGG motif [28]. The methylation status of the gene promoter regions can modulate the binding of RNA polymerase and/or transcription factors and can affect gene expression levels. DNA methylation patterns are modulated by growth conditions via changes in regulatory proteins that interact with DNA methylation sites [29]. Kahramanoglou *et al*. performed high-throughput sequencing of bisulfite-treated genomic DNA and microarray gene expression analysis, and they reported that DNA cytosine methylation regulates gene expression in *E. coli* stationary phase cells [28].

We demonstrated that the expression of the UV resistance genes *uvrA*, *uvrB*, *cho*, *umuC*, and *umuD* was strongly induced in stationary phase cells after UV irradiation (Fig. 4). We further demonstrated that the stationary phase cells of the dcm DNA cytosine methylase mutant exhibited high UV resistance compared to that of the wild-type strain (Fig. 5). Stationary phase-specific genes are further upregulated in the *dcm* strain [28]. We observed increased expression of *uvrB*, *cho*, *umuC*, *umuD*, and *sulA* in the stationary phase cells of the *dcm* strain compared to that in the wild-type strain (Fig. 6). This suggests that the DNA cytosine methylation state contributes to gene expression, ultimately resulting in a high UV resistance phenotype in the stationary phase *E. coli* cells.

## Acknowledgments

We are grateful to Prof. Hideto Miyake, Prof. Norihiro Nishimura, and Dr. Mina Okamura of Mie University for their cooperation in providing access to the UV-LED device. We also thank Dr. Tomohiro Itoh, Dr. Shuichi Karita, Ms. Haruka Fujimoto, and Mr. Norito Katsuo of Mie University for their technical support.

## Author contributions

**Shunsuke Ichikawa:** Conceptualization, Methodology, Investigation, Visualization, Writing - Original Draft, Supervision, Project administration, Funding acquisition. **Mika Okazaki:** Investigation, Validation.

## Abbreviations

CFU: Colony-forming unit
CPD: Cyclobutane pyrimidine dimers
DAPI: 4’,6-diamidino-2-phenylindole
Dcm: DNA cytosine methyltransferase
FITC: fluorescein isothiocyanate
NER: Nucleotide excision repair
PBS: Phosphate-buffered saline
UV-LED: Ultraviolet light-emitting diode

